# ASGAL: Aligning RNA-Seq Data to a Splicing Graph to Detect Novel Alternative Splicing Events

**DOI:** 10.1101/260372

**Authors:** Luca Denti, Raffaella Rizzi, Stefano Beretta, Gianluca Della Vedova, Marco Previtali, Paola Bonizzoni

## Abstract

**Background:** While the reconstruction of transcripts from a sample of RNA-Seq data is a computationally expensive and complicated task, the detection of splicing events from RNA-Seq data and a gene annotation is computationally feasible. The latter task, which is adequate for many transcriptome analyses, is usually achieved by aligning the reads to a reference genome, followed by comparing the alignments with a gene annotation, often implicitly represented by a graph: the *splicing graph*.

**Results:** We present `ASGAL` (Alternative Splicing Graph ALigner): a tool for mapping RNA-Seq data to the splicing graph, with the main goal of detecting novel alternative splicing events. `ASGAL` receives in input the annotated transcripts of a gene and an RNA-Seq sample, and it computes (1) the spliced alignments of each read, and (2) a list of novel events with respect to the gene annotation.

**Conclusions:** An experimental analysis shows that, by aligning reads directly to the splicing graph, `ASGAL` better predicts alternative splicing events when compared to tools requiring spliced alignments of the RNA-Seq data to a reference genome. To the best of our knowledge, `ASGAL` is the first tool that detects novel alternative splicing events by directly aligning reads to a splicing graph.

**Availability:** Source code, documentation, and data are available for download at http://asgal.algolab.eu.

## Background

Data coming from high throughput sequencing of RNA (RNA-Seq) can shed light on the diversity of transcripts that result from Alternative Splicing (AS). Three main computational approaches characterize transcriptome analysis from RNA-Seq data: (i) de novo assembly of transcript isoforms, (ii) gene annotation guided reconstruction of isoforms, and (iii) detection of AS events from a gene annotation or from an assembly graph. Various tools have been proposed in the literature that use the first two approaches. Examples of tools in category (i) that do not require a reference genome are `Trinity` [1] and `ABySS` [2]. `Cufflinks` [3], `Scripture` [4], and `Traph` [5], among many others, are known tools of category (ii). The first two tools were originally designed for de novo isoform prediction and can make limited use of existing annotations.

The final goal of reconstructing transcripts may be the detection of AS events that characterize gene expression. This step usually requires the comparison of a large number of transcripts that may arise from a sample of RNA-Seq reads. Such a comparison is performed by `AStalavista` [6], a popular tool for the exhaustive extraction and visualization of complex AS events from full-length transcripts. This tool does not use RNA-Seq reads as input but only the gene annotation, and it does not focus on single events (such as exon skipping, alternative splice sites, etc.) but rather uses a flexible coding of AS events [7] to list all the AS events between each pair of transcripts.

Since reconstructing full-length isoforms (either de novo or using a reference) from RNA-Seq reads is a difficult and computationally expensive problem, one may restrict the task to the direct detection of AS events from RNA-Seq data. Along this line of research, we propose a computational approach to predict AS events, and we implement this procedure in a tool belonging to category (iii). Similar tools are `SpliceGrapher` [8] and `SplAdder` [9] which take as input the spliced alignments of sequencing data (RNA-Seq data for `SplAdder`, and RNA-Seq data in addition to EST data for `SpliceGrapher`) against a reference genome, and produce as output a set of AS events. To do so, they exploit an augmented graph representation of the annotated transcripts, traditionally known as the *splicing graph* [10], with nodes and arcs that may represent novel AS events. In any case, while `SpliceGrapher` does not explicitly infer novel AS events, the main task of `SplAdder` is the prediction of AS events that are expressed by an input sample, and the quantification of those events by testing the differences between multiple samples. Two crucial computational instruments are usually required by tools of category (iii): an input file consisting of the alignment of RNA-Seq data to a reference genome, and the splicing graph. The first input may significantly change the performance of such tools, as the accuracy of the alignment may effect the predictions of AS events.

To avoid the possible bias due to the alignment against a reference genome, and motivated by the need of a direct comparison of RNA-Seq reads to the splicing graph, we propose `ASGAL` (Alternative Splicing Graph ALigner), a tool that consists of two parts: (i) a splice-aware aligner of RNA-Seq reads to a splicing graph, and (ii) a predictor of AS events supported by the RNA-Seq mappings. Currently, there are several tools for the spliced alignment of RNA-Seq reads against a reference genome or a collection of transcripts, but, to the best of our knowledge, `ASGAL` is the first tool specifically designed for mapping RNA-Seq data directly to a splicing graph. Differently from `SplAdder`, which enriches a splicing graph representing the gene annotation using the splicing information contained in the input spliced alignments, and then analyzes this enriched graph to detect the AS events differentially expressed in the input samples, `ASGAL` directly aligns the input sample to the splicing graph of the gene of interest and then detects the AS events which are novel with respect to the input gene annotation, comparing the obtained alignments with it. For this reason, `ASGAL` is designed to detect AS events also from an RNA-Seq sample, where the current annotation may differ by a single transcript per gene.

The approach of inferring AS events directly from RNA-Seq reads, without assembling isoforms, is also proposed in [11], where the main idea is to perform a de *novo* prediction of some AS events from the De Brujin graph assembly of RNA-Seq data, i.e. without using any gene annotation. An investigation of the de novo prediction of AS events directly from RNA-Seq data is also given in [12], where a characterization of the splicing graph that may be detected in absence of a gene annotation (either given as a reference or as a list of transcripts) is provided.

The `ASGAL` mapping algorithm improves a previous solution to the approximate pattern matching problem to a hypertext (an open problem faced in [13]). The approximate matching of a string to a graph with labeled vertices is a computational problem first introduced by Manber and Wu [14] and attacked by many researchers [15, 16, 17]. Navarro [18] improved all previous results in both time and space complexity, proposing an algorithm which requires 𝒪(*m*(*n* + *e*)) time, where m is the length of the pattern, n is the length of the concatenation of all vertex labels, and e is the total number of edges. The method in [13] improves the latest result by Thachuk [19]: an algorithm with time complexity 𝒪 (*m* + γ^2^) using succinct data structures to solve the exact version of matching a pattern to a graph — i.e. without errors — where γ is number of occurrences of the node texts as substrings of the pattern. The algorithm in [13] is based on the concept of *Maximal Exact Match* and it uses a succinct data structure to solve the approximate matching of a pattern to a hypertext in 𝒪 (*m* + *η*^2^) time, where *η* is the number of Maximal Exact Matches between the pattern and the concatenation of all vertex labels. In this paper we improve the results in [13] by extending the algorithm to implement an RNA-Seq data aligner for detecting general AS events from the splicing graph.

An experimental analysis on real and simulated data was performed with the purpose of assessing the quality of `ASGAL` in detecting AS events. We compared `ASGAL` with `SplAdder`, this latter using both `Hisat2` [20] (the successor of `TopHat2` [21]) and `STAR` [22] as spliced aligners. Since `ASGAL` aligns reads directly to the splicing graph, it reveals a higher precision in predicting AS events. Although `ASGAL` works under different assumptions than other existing tools, we decided to compare `ASGAL` with `SplAdder`, since in [9] the authors showed that `SplAdder` achieves better re-sults than other similar tools such as `rMATS` [23] and `JuncBASE` [24]. In particular, `rMATS` and `JuncBASE` are tools that only detect AS events from the differential expression of transcripts in multiple samples of RNA-Seq data. To do this, `JuncBASE` uses a complex pipeline containing other tools such as `Cufflinks` and requires long running times, while `rMATS` has limited capacity to infer AS event from RNA-Seq data alignments [9].

## Methods

`ASGAL` (Alternative Splicing Graph ALigner) is a tool for performing a mapping of RNA-Seq data in a sample against the splicing graph of a gene with the main goal of detecting the novel splicing events expressed by the reads of the sample with respect to the annotation of the gene. More precisely, `ASGAL` takes as input the annotation of a gene together with the related reference sequence, and a set of RNA-Seq reads, to output (i) the spliced alignments of each read in the sample and (ii) the alternative splicing events expressed in the sample which are novel with respect to the annotation. We point out that `ASGAL` uses the input reference sequence mainly for building the splicing graph. Each identified event is described by its type, *i.e.* exon skipping, intron retention, alternative acceptor splice site, alternative donor splice site, its genomic positions, and a measure of its quantification, *i.e.* the number of reads that support it.

This section is organized as follows. We first introduce the basic definitions and notions that we will use in the section *spliced graph-alignment*, and finally we describe the steps of our method.

### Definitions

From a computational point of view, a genome is a sequence of characters, *i.e.* a string, drawn from an alphabet of size 4 (`A, C, G`, and `T`). A gene is a *locus* of the genome, that is, a gene is a substring of the genome. Exons and introns of a gene locus will be uniquely identified by their starting and ending positions on the genome. A transcript T of gene G is a sequence ⟨[a_1_, b_1_], [a_2_, b_2_],…, [a_n_, b_n_]⟩ of exons on the genome, where a_*i*_ and b_*i*_ are respectively the *start* and the *end* positions of the *i*-th exon of the transcript. Observe that *a*_1_ and *b*_n_ are the *starting* and *ending positions* of transcript *T* on the genome, and each [*b*_i_ + 1, *a*_*i*+1_ — 1] is an *intron* represented as a pair of positions on the genome. In the following, we denote by *ε*_*G*_ the set of all the exons of the transcripts of gene *G*, that is 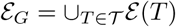, where *ε* (*T*) is the set of exons of transcript *T* and 𝒯 is the set of transcripts of *G*, called the *annotation* of *G*. Given two exons *e*_*i*_ = [*a_i_,b_i_*] and e_*j*_ = [*a_j_,b_j_*] of *ε*_G_, we say that *e*_*i*_ *precedes e*_*j*_ if *b*_*i*_ < *a*_*j*_ and we denote this by *e*_*i*_ ≺ e_*j*_. Moreover, we say that *e*_*i*_ and *e*_*j*_ are *consecutive* if there exists a transcript *T* ∈ 𝒯 and an index *k* such that e_*k*_ = *e*_*i*_ and *e*_*k*+1_ = *e*_*j*_.

The *splicing graph* of a gene *G* is the directed acyclic graph *S*_*G*_ = (*ε*_*G*_, *E*), *i.e.* the vertex set is the set of the exons of *G*, and the edge set *E* is the set of pairs (*v*_*i*_, *v*_*j*_) such that *v*_*i*_ and *v*_*j*_ are consecutive. For each vertex *v*, we denote by seq(*v*), the genomic sequence of the exon associated to *v*. Finally, we say that 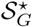 is the graph obtained by adding to S_G_ all the edges (*v*_*i*_, *v*_*j*_) ∉ *E* such that *v*_*i*_, ≺ *v*_*j*_. We call these edges *novel* edges. Note that the novel edges represent putative junctions between two exons that are not consecutive in any transcript of *G* and will be used to detect novel alternative splicing events of *G* induced by a set of RNA-Seq reads.

In the following, we will use the notion of *Maximal Exact Match* (MEM) to perform the spliced graph-alignment of an RNA-Seq read to *S*_*G*_. Given the two strings *H* and *R*, a MEM is a triple *m* = (*i*_H_, *i*_R_, *l*) representing the common substring of length *l* between the two strings that starts at position *i*_H_ in *H*, at position *i*_R_ in *R*, and that cannot be extended in either direction without introducing a mismatch. Computing the MEMs between a string *R* and a splicing graph *S*_*G*_ can be done by concatenating the labels of all the vertices and interposing the special symbol φ, obtaining a string *H* that we call the *linearization* of the splicing graph. Then, by employing the algorithm by Ohlebusch *et al* [25], all the MEMs between R and *S*_*G*_ can be computed in linear time with respect to the length of the reads and the number of MEMs. It is immediate to see that, given a vertex *v* of *S*_*G*_, the label seq(*v*) is a substring *H*[*i*_*v*_, *j*_*v*_] of the linearization of *S*_*G*_ and a MEM must occur inside a vertex label. In the following, given a read *R* and the linearization *H* of *S*_*G*_, we say that a MEM *m* = (*i*_*H*_, *i*_*R*_, *l*) occurs inside the vertex label seq(*v*) if *i*_H_ is a position of the interval [*i*_*v*_,*j*_*v*_]. We say that a MEM *m* = (*i*_*H*_, *i*_*R*_,*l*) precedes another MEM 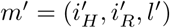 in *R* if 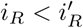 and 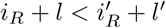, and we denote this by *m* ≺ _*R*_ *m*′. Similarly, when *m* precedes *m*′ in *H*, we denote it by *m* ≺_*H*_ *m′*, if the previous properties hold and the two MEMs belong to the same vertex label `seq`(*v*). When m precedes m′ in *R* (in *H*, respectively), we say that 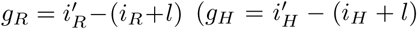, respectively) is the gap between the two MEMs. If the gaps *g*_*R*_ or *g*_*H*_ (or both) are positive, we refer to the gap strings as *G*_*R*_ and *G*_*H*_, while when they are negative, we say that *m* and *m*′ *overlap* either in *R* or *H* (or both). Given a MEM *m* belonging to the vertex labeled `seq`(*v*), we denote as `PREF`_*H*_(*m*) and `SUFF`_*H*_(*m*) the prefix and the suffix of `seq`(*v*) upstream and downstream from the start and the end of *m*, respectively.

### Spliced graph-alignment

We are now able to define the fundamental concepts that will be used in our method. In particular, we first define a general notion of gap graph-alignment and then we introduce specific constraints on the use of gaps to formalize a splice-aware graph-alignment that is fundamental for the detection of alternative splicing events in `ASGAL`.

A *gap graph-alignment* of *R* to graph *S*_*G*_ is a pair (*A*,*π*) where *π* = ⟨*v*_1_,…, *v*_*k*_⟩ is a path of the graph 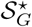and

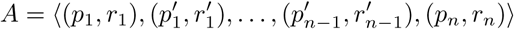

is a sequence of pairs of strings such that `seq`(*v*_1_) = *x* · *p*_1_ and `seq`(*v*_*n*_) = *p*_*n*_ · *y*, for *x, y* eventually empty strings and 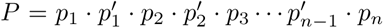 is the string labeling the path *π* and 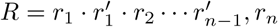

The pair (*p_i_,r_i_*), called a *factor* of the alignment *A*, consists of a non-empty substring *r*_*i*_ of *R* and a non-empty substring *p*_*i*_ of the label of a vertex in *π*. The pair 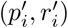, called a *gap-factor* of the alignment *A*, consists of at least an empty substring ϵ. Moreover, either 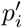 is empty or 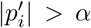, and either 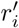 is empty or 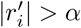, for a fixed value α.

We associate to each factor (*p_i_,r_i_*) the cost δ(*p_i_,r_i_*), and to each gap-factor 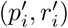 the cost 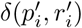, by using a function *δ*(·,·) with positive values. Then the cost of the alignment (*A, π*) is given by the expression:

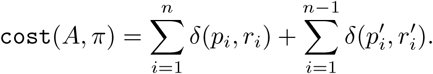

Notice that the constraint on the length of strings in a gap-factor derives from the intuition that we want gap-factors to represent events induced by a gap of a given length. Moreover, we define the *error* of a gap graph-alignment as the sum of the edit distance of each factor (but not of gap-factors). Formally, the error of the alignment ( *A, π*) is:

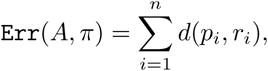

where *d*(·, ·) is the edit distance between two strings. Intuitively, in a gap graph-alignment, factors correspond to portions of exons covered by portions of the read, while gap-factors correspond to new splice junctions or new splicing events induced by the alignment of the read. To define a splice-aware alignment, that we call *spliced graph-alignment*, we need to classify each gap-factor and to assign it a cost. Our primary goal is to compute a gap graph-alignment of the read to the splicing graph that possibly reconciles to the gene annotation; if this is not possible, then we want to minimize the number of novel events. For this reason we distinguish three types of gap-factors: *annotated, novel*, and *impossible*. Intuitively, an annotated gap-factor models an annotated intron, a novel gap-factor represents a novel intron which can represent an alternative splicing event, while an impossible gap-factor does not represent any intron.

Formally, we classify a gap-factor 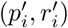 as *annotated* if and only if 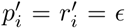 and the two strings *p*_*i*_, *p*_i+1_ are on two different vertices that are linked by an edge in *S*_*G*_.

We classify a gap-factor 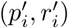 as novel in the following cases:

1. 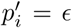 occurs between the strings *p*_*i*_ and *p*_i+1_ which belong to two distinct vertices linked by an edge in 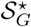 *(i.e.* this gap-factor represents an exon skipping, an alternative splicing site or a cassette exon event - Figure 1, cases *a* and *d*).
2. 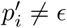 occurs between the strings *p*_*i*_ and *p*_*i*+1_ which belong to the same vertex of 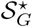 *(i.e.* this gap-factor represents an intron retention event - Figure 1, case b).
3. 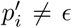 occurs between the strings *p*_*i*_ and *p*_i+1_ which belong to two distinct vertices linked by an edge in 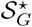 *(i.e.* this gap-factor represents an alternative splice site event - Figure 1, case *c*).

**Figure 1.**
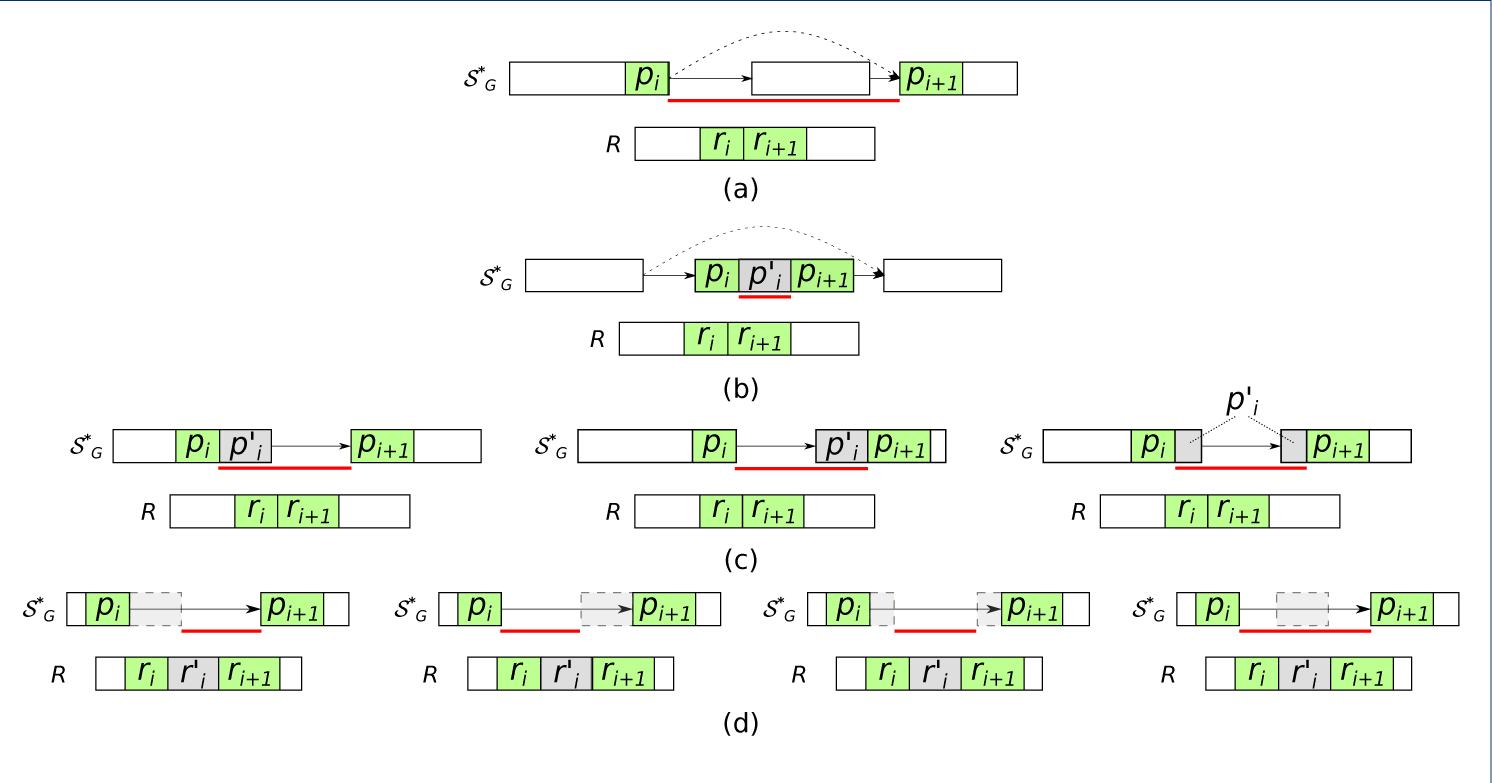
Relationship between novel gap-factors, introns, and AS events. All the possible gap-factors 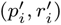 that can occur in a spliced graph-alignment are shown. Each subfigure depicts a splicing graph (actually, 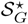), a read R and a portion of a possible spliced graph-alignment: the green squares represent the factors while the gray ones represent the gap-factors. The red line below each splicing graph, instead, represents the intron supported by the alignment that is used to infer the possible presence of alternative splicing events. In more detail, case (a) shows an exon skipping, case (b) shows an intron retention, case (c) shows alternative splice sites shortening an exon, and case (d) shows alternative splice sites extending an exon or a cassette exon.

In the remaining cases, which are (i) 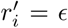 and 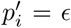 occurs between strings *p*_*i*_ and *p*_*i*+1_ which belong to the same vertex, and (ii) 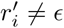 and 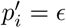 occurs between strings *p*_*i*_ and *p*_*i*+1_ which belong to the same vertex, the gap-factor is classified as impossible. We notice that in the former case, factors (*p*_*i*_,*r*_*i*_) and (*p*_*i*+1_,*r*_*i*+1_) can be joined into an unique factor.

Let 𝒢_𝓕_ be the set of novel gap-factors of a gap graph-alignment *A*. Then a *spliced graph-alignment* (*A, π*) of *R* to *S*_*G*_ is a gap graph-alignment in which impossible gap-factors are not allowed, whose cost is defined as the number of novel gap-factors, and whose error is at most *β* — for a given constant *β*. In other words, in a spliced graph-alignment (*A, π*), we cannot have impossible gap-factors, and the *δ* function assigns a cost 1 to each novel gap-factor and a cost 0 to all other factors: thus `cost`(*A, π*) = |𝒢_𝓕_| and `Err`(*A, π*) ≤*β*. We focus on a bi-criteria version of the computational problem of computing the optimal *spliced graph-alignment* (*A, π*) of *R* to a graph *S*_*G*_, where first we minimize the cost, then we minimize the error. The intuition is that we want a spliced graph-alignment of a read that is consistent with the fewest novel splicing events that are not in the annotation. Moreover, among all such alignments we look for the alignment that has the smallest edit distance (which is likely due to sequencing errors) in the non-empty regions that are aligned (*i.e.* the factors). Figure 2 shows an example of spliced graph-alignment of error value 2, and cost 2 — since it has two novel gap-factors.

**Figure 2.**
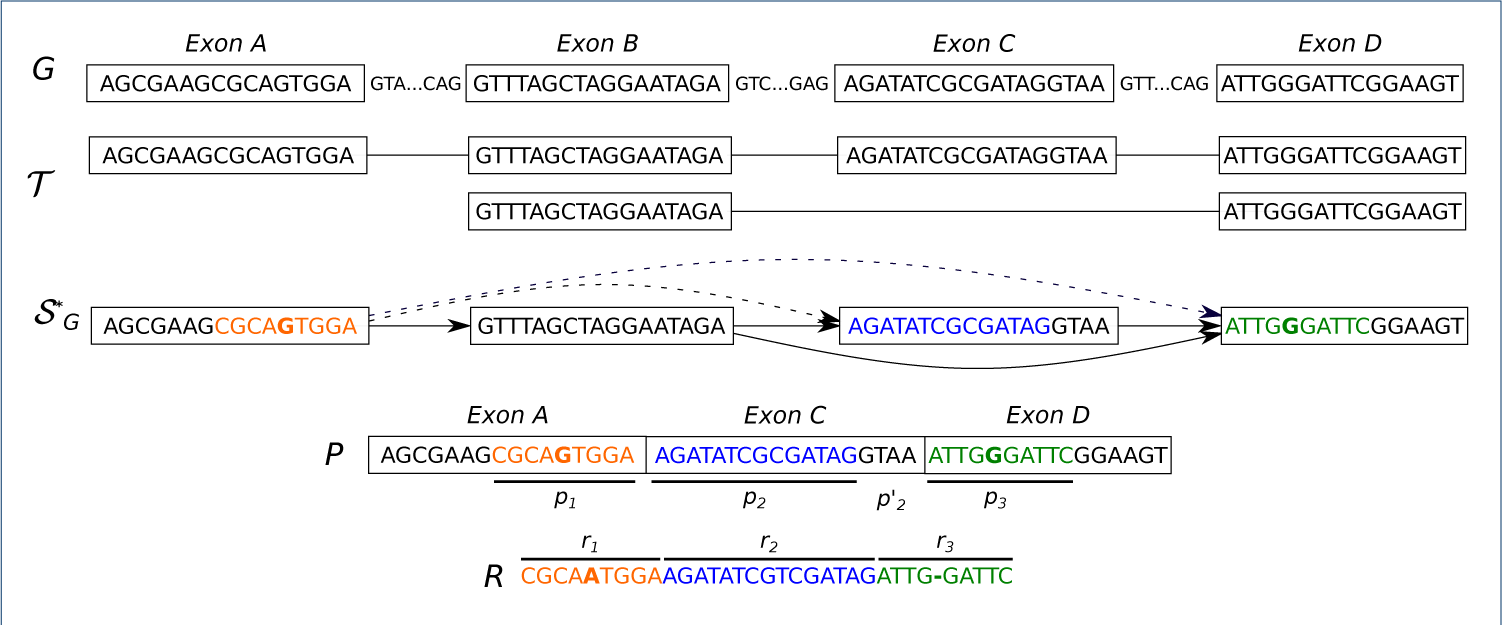
Spliced graph-alignment. Example of *spliced graph-alignment* of a read *R* to the splicing graph of a simple gene with four exons *(A, B, C*, and *D*) and two transcripts. The splicing graph 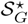 is depicted where dashed edges represent the novel edges. The read *R* has been factorized in three strings *r*_*1*_, *r*_2_ and *r*_3_ matching to strings *p*_1_, *p*_2_ and *p*_3_ of *P* (which is the concatenation of exon labels of path *π* = ⟨A,C, D⟩). We observe that 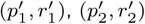 are two novel gap-factors, *r*_*1*_ matches *p*_1_ with an error of substitution while *r*_3_ matches *P*_3_ with an error of insertion: both the error and the cost of this spliced-graph alignment are equal to 2. This alignment of *R* to the splicing graph of *G* supports the evidence of two *novel* alternative splicing events: the skipping of exon *B* and the alternative donor site of exon *C*.

In this paper we propose an algorithm that, given a read *R*, a splicing graph S_G_, two constants *α* and *β*, computes an optimal spliced graph-alignment — that is, a spliced graph-alignment with minimum cost and, among all spliced graph-alignments with minimum cost, we compute the alignment with minimum error. Once the spliced graph-alignments are computed, the novel gap-factors are compared with the reference genome to determine which novel AS events are induced.

The next section details how `ASGAL` computes the spliced graph-alignments of a set of reads to the splicing graph *S*_*G*_, how it selects an optimal spliced graph-alignment, and how it exploits novel gap-factors to detect AS events.

### ASGAL approach

We now describe the algorithm employed by `ASGAL` to compute optimal spliced graph-alignments, *i.e.* by aligning the RNA-Seq reads of a sample to the splicing graph of a gene. Then alignments to the splicing graph are used to identify the Alternative Splicing events. The `ASGAL` tool implements a pipeline consisting of the following steps: (1) construction of the splicing graph of the gene, (2) computation of the spliced graph-alignments of the RNA-Seq reads, (3) translation of the alignments from the splicing graph to the genome, and (4) detection of the novel alternative splicing events.

In the first step, `ASGAL` builds the splicing graph S_G_ of the input gene using the reference genome and the gene annotation, and adds the novel edges to obtain the graph SG which will be used in the next steps.

The second step of `ASGAL` computes the spliced graph-alignments of each read R in the input RNA-Seq sample by combining MEMs in factors and gap-factors. To do so, we extend the approximate pattern matching algorithm of Beretta *et al.* [13] to obtain the spliced graph-alignments of the reads, which will be used in the following steps to detect novel alternative splicing events.

As anticipated before, we use the approach proposed by Ohlebusch *et al.* in [25] to compute, for each input read *R*, the set of MEMs between *H*, the linearization of the splicing graph *S*_*G*_, and *R* with minimum length *L*, a user-defined parameter (we note that the approach of [25] allows to specify the minimum length of MEMs). We recall that the string H is obtained by concatenating the strings `seq`(*v*) and φ for each vertex *v* of the splicing graph. We point out that the concatenation order does not affect the resulting alignment and that the splicing graph linearization is performed only once before aligning the input reads to the splicing graph.

Once the set *M* of MEMs between *R* and *H* is computed, we build a weighted graph *G*_*m*_ = (*M, E*_*M*_) based on a given parameter *α* and the two precedence relations between MEMs, ≺_*R*_ and ≺_*H*_, respectively. Then we use such graph to extract factors and gap-factors. More precisely, the vertex set is the set *M* of MEMs between *H* and *R*, and there exists an edge from *m* to *m*′, with *m, m*′ *∈ M*, if and only if *m* ≺_R_ *m*′ and one of the following conditions, also depicted in Figure 3, holds:

1. *m* and *m*′ are inside the same vertex label of *H, m* ≺_*H*_ *m*′, and either (i) *g*_*R*_ > 0 and *g*_*H*_ > 0, or (ii) *g*_*R*_ = 0 and 0 < *g*_*H*_ ≤ *α*. The weight of the edge (*m, m*′) is set to the edit distance between *G*_*R*_ and *G*_*H*_.
2. *m* and *m*′ are inside the same vertex label of *H, m* ≺_*H*_ *m*′, *g*_*R*_ ≤ 0, and g_H_ ≤ 0. The weight of the edge (*m, m*′) is set to |*g*_*R*_ - *g*_*H*_|.
3. *m* and *m*′ are inside the same vertex label of *H, m* ≺_*H*_ *m*′, *g*_*R*_ ≤ 0 and *g*_*H*_ > α. The weight of the edge (*m, m*′) is set to 0.
4. *m* and *m*′ are on two different vertex labels `seq`(*v*_1_) and `seq`(*v*_2_), with *v*_1_ ≺ *v*_2_, and *g*_*R*_ ≤ 0. The weight of the edge (*m, m*′) is set to 0.
5. *m* and *m*′ are on two different vertex labels `seq`(*v*_1_) and `seq`(*v*_2_), with *v*_1_ ≺ *v*_2_, *g*_*R*_ > 0, and `SUFF`_*H*_(*m*) = `PREF`_*H*_(*m*′) =?. The weight of the edge (*m, m*′) is set to 0 if *g*_*R*_ > α, and to *g*_*R*_ otherwise.
6. *m* and *m*′ are on two different vertex labels seq(*v*_1_) and seq(*v*_2_), with *v*_1_ ≺ *v*_2_,*g*_*R*_ > 0, at least one between `SUFF`_*H*_(*m*) and `PREF`_*H*_(*m*′) is not ϵ. The weight of the edge (*m, m*′) is set to the edit distance between *G*_*R*_ and the concatenation of `SUFF`_*H*_(*m*) and `PREF`_*H*_(*m*′).

**Figure 3.**
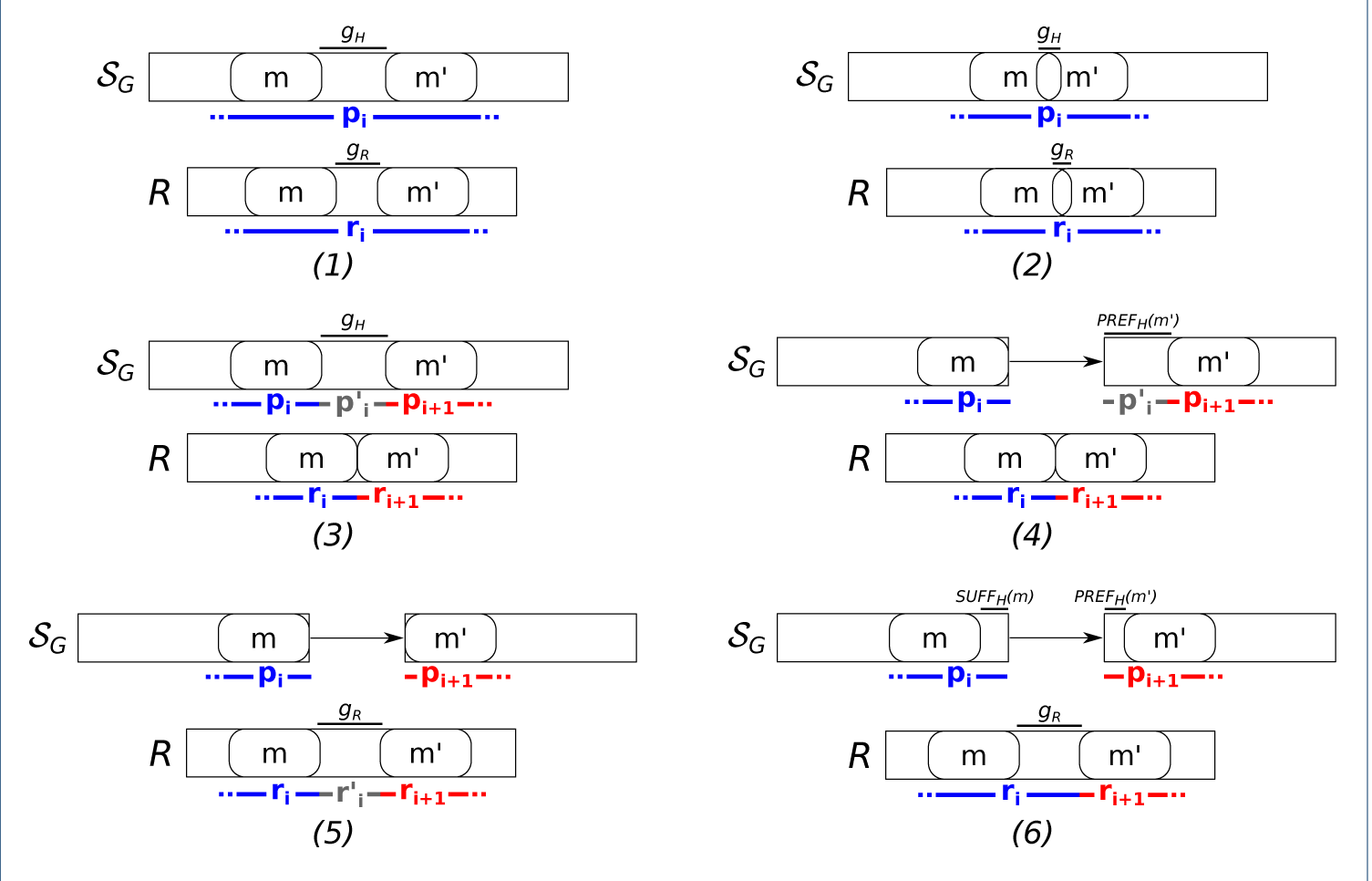
Conditions for linking two different MEMs All the conditions used to connect two different MEMs and then to build the factors and gap-factors of a spliced graph-alignment are shown. In all the conditions, the first MEM must precede the second one on the read. In condition (1) and (2), the two MEMs occur inside the same vertex label and leave a gap (condition 1) or overlap (condition 2) on the read or on the vertex label. In these conditions, the two MEMs are joined in the same factor of the alignment. In condition 3, instead, the two MEMs occur inside the same vertex label but they leave a long gap only on the vertex label and not on the read. In this case, the two MEMs belong to two different factors linked by a gap-factor. In the other conditions, instead, the two MEMs are inside the labels of two different vertices of the splicing graph, linked by a (possible novel) edge. For this reason, in any of these cases, the two MEMs belong to two different factors of the alignment. In condition 4, the two MEMs leave a gap only the path, in condition 5 they leave a gap only on the read, and in condition 6, they leave a gap on both the path and the read.

Notice that the aforementioned conditions do not cover all of the possible situations that can occur between two MEMs, but they represent those that are relevant for computing the spliced graph-alignments of the considered read. Intuitively, conditions 1 and 2 are used to handle the possible presence of alignment errors, conditions 3, 4, and 5 are used to model annotated or novel gap-factors of the alignment, and condition 6 is used for both these purposes.

The spliced graph-alignment of each read *R* is computed by a visit of the graph *G*_*m*_. More precisely, each path *π*_*Μ*_ of this graph represents a spliced graph-alignment and the weight of the path is the number of differences between the pair of strings in *R* and *H* covered by *π*_*Μ*_. For this reason, for read *R*, we select the lightest path in *G*_*M*_, with weight less than *β* (a given error threshold) which also contains the minimum number of novel gap-factors, *i.e.* we select an optimal spliced graph-alignment. In detail, given an edge (*m, m*′), if either condition 1 holds or condition holds, *m* and *m*′ are candidates to be merged inside a factor. If condition 3 holds, then *m* and *m*′ are candidates to be located respectively in the suffix and the prefix of two consecutive factors of the alignment which are separated by the novel gap-factor 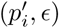, with 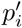 representing the string between the two MEMs in 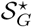 (gap-factor of type 2 in the classification of novel gap-factors given in Section *Spliced graph-alignment*). If one of the remaining conditions 4, 5, 6 holds, then *m* and *m*^′^ are candidates to be located respectively in the suffix and in the prefix of two consecutive factors (*p*_*i*_,*r*_*i*_) and (*p*_i+1_,*r*_i+1_) of the alignment separated by the gap-factor 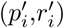. Moreover, if condition 4 holds, then 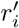 is ϵ and 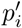 is the string between the two MEMs in 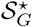. Then, if 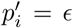, then 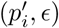 is a novel gap-factor of type 1 or an annotated gap-factor; otherwise, it is a novel gap-factor of type 3. If condition 5 holds, then 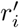 is the string between the two MEMs in *R* and 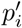 is ϵ. In this case, 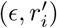 is a novel gap-factor of type 1 or an annotated gap factor. Finally, if condition 6 holds, then both 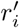 and 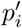 are ϵ and they identify either a novel gap-factor of type 1 or an annotated gap-factor.

The third step of `ASGAL` computes the spliced alignments of each input read with respect to the reference genome starting from the spliced graph-alignments computed in the previous step. Exploiting the annotation of the gene, we convert the coordinates of factors and gap-factors in the spliced graph-alignment to positions on the reference genome. In fact, observe that factors map to coding regions of the genome whereas gap-factors identify the skipped regions of the reference, *i.e.* the introns induced by the alignment, modeling the possible presence of AS events (see Figure 1 for details). We note here that converting the coordinates of factors and gap-factors to positions on the reference genome is pretty trivial except when factors *p*_*i*_ and *p*_*i*+1_ are on two different vertices and only 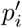 is ϵ (case *d* of Figure 1). In this case, the portion 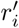 must be aligned to the intron between the two exons whose labels contains *p*_*i*_ and *p*_*i*+1_ as a suffix and prefix, respectively. If 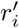 aligns to a prefix or a suffix of this intron (taking into account possible errors within the total error bound α), then the left or right coordinate of the examined intron is modified according to the length of 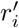 (first and second cases of Figure 1(d)). In the other cases (third and fourth cases of Figure 1(d)), the portion 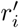 is not aligned to the intron and it is represented as an insertion in the alignment. Notice that in these latter cases, 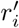 may support either a novel exon on the genome, two alternative splice sites or even a more complex combination of these events. Moreover, the third step of our approach performs a further refinement of the splice sites of the introns in the obtained spliced alignment since it searches for the splice sites (in a maximum range of 3 bases with respect to the original ones) determining the best intron pattern (firstly *GT-AG*, secondly *GC-AG* if *GT-AG* has not been found).

In the fourth step, `ASGAL` uses the set 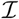 of introns supported by the spliced alignments computed in the previous step, *i.e.* the set of introns associated to each pair of gap-factors, to detect the alternative splicing events expressed in the given RNA-Seq sample with respect to the given annotation. Let 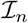 be the subset of 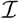 composed of the introns which are not present in the annotation, that is, the *novel* introns. For each *novel* intron 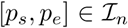 with at least ω supporting alignments, `ASGAL` identifies one of the following events:

−*exon skipping*, if there exists an annotated transcript containing two non-consecutive exons [*a*_*i*_,*p*_*s*_ — 1] and [*p*_*e*_ + 1, *b*_*j*_].
−*intron retention*, if there exists an annotated transcript containing an exon [*a*_*i*_, *b*_*i*_] such that (i) *a*_*i*_ < *p*_*s*_ < *p*_*e*_ < *b*_*i*_, (ii) there exists an intron in 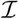 ending at *a*_*i*_ — 1 or *a*_*i*_ is the start of the transcript and (iii) there exists another intron in 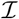 starting at *b*_*i*_ + 1 or *b*_*i*_ is the end of the transcript.
−*alternative acceptor site*, if there exists an annotated transcript containing two consecutive exons [*a*_*i*_,*p*_*s*_ — 1] and [*a*_*j*_, *b*_*j*_] such that *p*_*e*_ < *b*_*j*_, and there exists an intron in 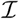starting at *b*_*j*_ + 1 or *b*_*j*_ is the end of the transcript.
−*alternative donor site*, if there exists an annotated transcript containing two consecutive exons [*a*_*i*_, *b*_*i*_] and [*p*_*e*_ + 1, *b*_*j*_] such that *p*_*s*_ > *a*_*i*_, and there exists an intron in 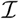ending at *a*_*i*_ — 1 or *a*_*i*_ is the start of the transcript.

## Results

In this section we will describe the experimental evaluation we performed to check the ability of `ASGAL` to align the reads to the splicing graph and to detect alternative splicing events. Such experimental analysis was done on both simulated and real data, in which the former had the specific goal of measuring the quality of our tool, whereas the latter of proving the ability of `ASGAL` in dealing with real datasets. In all our experiments, we have run `ASGAL` with its default values: the minimum length of the MEMs L is 15, *α* and *β* is 3% of the maximum length of an input read, and the minimum support for AS events *ω* is 3.

### Simulated Data

In the first phase of our experimental analysis, we evaluated our tool using simulated data. The goal of this analysis was twofold: to assess the efficiency of our method in aligning a set of RNA-Seq reads against a splicing graph, and to assess the usefulness of the alignment method to detect the alternative splicing events expressed by the sample with respect to a given annotation.

Since the alignment steps of `ASGAL` were tailored to the identification of novel alternative splicing events, the quality of the alignments was not directly assessed by a comparison with other spliced aligners. In fact, a first part of the experimental analysis measures how often each read is mapped to the correct gene by our approach, `STAR` [22], and `Hisat2` [20] (the successor of `TopHat2` [21], another well-known spliced aligner).

Then, in a second step we compared `ASGAL` with `SplAdder` [9] to assess their accuracy in identifying alternative splicing events. To avoid any bias in the experiments, we decided to use the same reference genome, annotations and simulated data^[1]^ used in [9]. From these data, simulated using `Flux` [26], we considered two different RNA-Seq datasets with 5 million and 10 million reads respectively, each covering 1000 genes randomly selected from the human GENCODE annotation (v19) [27]. Then, we divided each dataset into 24 samples, that is one for each chromosome, and we used `cutadapt` [28] to remove poly-A tails.

Since our tool works at gene level — that is, it considers the splicing graph of each gene independently — to perform a fair comparison, in the first step of our experiments on simulated data, we ran `STAR` and `Hisat2` on the exact genomic region of each gene, cut from the reference genome based on the given annotation. We ran all tools using a single thread, to measure the time performance, and providing the default parameters. The only exception to the latter condition is that the `genomeSAindexNbases` option of `STAR` was set according to the length of the genomic regions^[2]^ to avoid crashes during the alignment step: the default values of `STAR` are suitable only for longer reference sequences than what we used in our experiments.

Table 1 summarizes the precision, recall, and F-measure of the three tools on the dataset composed of 5M reads. These quality measures were computed for each gene by considering the number of reads simulated from the gene and aligned to that gene (true positives), the number of reads simulated from other genes and aligned to that gene (false positives), and the number of reads simulated from that gene and not aligned to it (false negatives). In this evaluation, we considered only the primary alignments of each read provided by the tested tool.

**Table 1.**
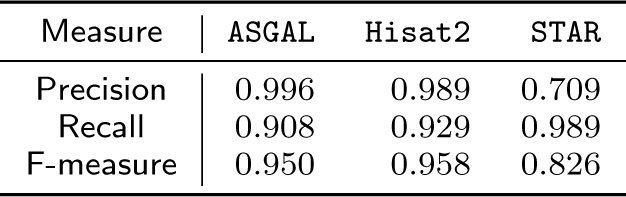
Q uality results of the alignments on the simulated datasets composed of 5M reads. For each of the tested methods, that is, `ASGAL, Hisat2`, and `STAR`, the values of Precision, Recall, and F-measure achieved in the alignment step on the simulated dataset are reported.

Our comparison of the aligners (Table 1) shows that `ASGAL` has the best precision and the worst recall, while `STAR` has the worst precision and the best recall. Moreover, `Hisat2` has the best F-measure, and `ASGAL` is within 1% of that. This confirms the ability of the method we propose to align a read to the correct gene, which was the main goal of this task. Notice that `ASGAL` considers only exonic regions encoded in a splicing graphs, while both `Hisat2` and `STAR` align against a genomic region.

For this reason, these two latter tools align reads also in intronic regions. Moreover, due to error rate (set to 3%) and the read length (100bp), `ASGAL` computes only primary alignments with maximum 3 errors while the other tools, by default, accept also primary alignments with a higher number of errors. This is the main reason for the worse recall of `ASGAL`.

Table 2 summarizes the computational resources (total time and memory peak) required in the alignment step by the considered tools. Analyses were performed on a 64 bit Linux (Kernel 4.4.0) system equipped with Four 8-core Intel Xeon 2.30GHz processors and 256GB of RAM. The time is the sum of the times required to compute the index of the input reference (the genomic sequence for `Hisat2` and `STAR`, the splicing graph for `ASGAL`) and then to align the reads.

**Table 2.** Computational resources required by the three tested methods (`ASGAL, Hisat2`, and `STAR`) to align the simulated datasets composed of 5M reads. These results are shown in terms of time (minutes) and memory peak (MegaBytes).

| Measure | ASGAL | Hisat2 | STAR |
| --- | --- | --- | --- |
| Time (m) | 763 | 70 | 9424 |
| Memory Peak (MB) | 267 | 98 | 275 |

While the memory peak usage is below 300MB for all tools, the running time varies greatly, with `STAR` using more than 100× the running time of `Hisat2` and `ASGAL` using more than 10× the running time of `Hisat2`. Notice that each gene was processed independently, hence the execution of the pipelines is embarrassingly parallel and we can fully use all available cores. Moreover, `STAR` is tailored for aligning against a complete reference genome, not against a set of much smaller genomic regions. For these reasons, we are not interested in comparing the computational efficiency of the tools, but only in assessing the feasibility of the approach.

The main goal of our experimental analysis on simulated data was to evaluate the capability of our tool to detect alternative splicing events. We decided to compare our tool with `SplAdder`, a software for identifying and quantifying alternative splicing events starting from a given annotation and the alignment files, since it is the best approach for detecting alternative splicing events, from the results obtained in [9].

Before comparing the results obtained by `ASGAL` with those computed by `SplAdder`, we would like to highlight that these two tools, although they could look similar, perform different tasks. More precisely, as proved by our experiments, `SplAdder` builds a splicing graph starting from a given annotation and enriches it by exploiting the spliced alignments, but to identify the AS events it requires that all the isoforms involved in the event are supported by reads in the sample. On the other hand, `ASGAL` identifies the alternative splicing events even if the sample contains only reads extracted from a single isoform, since it uses the annotation as a reference for the identification of the novel AS events. This case is especially important, since usually there is a single transcript expressed per gene, when considering a single sample [29]. As said in `SplAdder`’s supplemental material, the default behaviour of `SplAdder` can be modified by adapting different parameters that guide the confirmation process of each alternative splicing event found. However, it is not an easy task to modify these parameters since they are hard-coded and it is not even clear how to choose the best values without the risk of introducing undesired behaviors.

As done in [9], to assess the accuracy of the two tools, we provided them a reduced annotation, obtained in the following way. First of all, we used `AStalavista` [6] to extract all the alternative splicing events contained in the annotations of the 1000 considered genes. This resulted in a total of 2568 alternative splicing events: 1574 exon skippings, 416 alternative acceptor sites, 290 alternative donor sites, and 288 intron retentions. Then, for each gene and for each event identified by `AStalavista`, we created a new reduced annotation containing all the transcripts except those responsible for such event. Here, we focused our attention on exon skippings, alternative splice sites (both acceptor and donor), and intron retentions caused by the insertion of a new intron inside an exon. For completeness, we notice that we doubled the alternative splice sites events in order to test the considered tools in the detection of both alternative splicing sites events shortening and extending an exon while we did not consider in this experiments the possible insertion of a new exon inside an intron (cassette exon) and the intron retentions caused by the union of two exons. Moreover, when different events on the same gene produced the same reduced annotation, we considered the annotation only once. We obtained a total of 3274 AS events and 2792 reduced annotations.

We did not replicate exactly the experiments done in [9] — where the reduced annotation contained only the first transcript of each gene — since it would not have been a fair comparison. In fact, our tool heavily depends on the alignments of RNA-Seq reads against the splicing graph, while `SplAdder` takes as input the alignments against the full genome reference sequence.

For each gene and for each reduced annotation, we run the two tools and we evaluated their accuracy in identifying the events found by comparing the reduced and the full annotations. We have used two datasets of reads, respectively with 5M and 10M reads. For each kind of alternative splicing event we analyzed the predictions over the set of 1000 genes, computing the corresponding values of precision, recall, and F-measure. More precisely, given a gene and its reduced annotation, we consider as ground truth the set of events found by `AStalavista` in the original annotation, but not in the reduced annotation. To compute the values of precision, recall, and F-measure, we considered the number of events inducing the reduced annotation and found by the tool (true positives), the number of events inducing the reduced annotation not found by the tool (false negatives), and the number of events found not contained in the output of `AStalavista` for that gene (false positives). As anticipated, for each gene it is possible to have more events expressed in the reduced annotation than the wanted ones: these cases correspond to annotations in which the removal of the annotated transcript generated more than a single alternative splicing event. This offered the possibility to evaluate the behavior of `ASGAL` in dealing with complex scenarios. Table 3 summarizes the number of reduced annotations containing 1, 2, 3, 4, and 5 different alternative splicing events and how many events of each type they contain.

**Table 3.** Information about the reduced annotations considered in the experiments on simulated data. Each row summarizes the number of reduced annotation (#Annotations) containing the corresponding number of AS events (#Events), the total number of AS events (#TotEvents) and their partitioning in exon skipping (ES), alternative acceptor site (A3), alternative donor site (A5), intron retention (IR). The last row shows the sum of the values contained in each column.

| #Events | #Annotations | #TotEvents | ES | A3 | A5 | IR |
| --- | --- | --- | --- | --- | --- | --- |
| 1 | 2393 | 2393 | 1341 (56%) | 519 (22%) | 380 (16%) | 153 (6%) |
| 2 | 329 | 658 | 205 (31%) | 208 (32%) | 146 (22%) | 99 (15%) |
| 3 | 58 | 174 | 26 (15%) | 70 (40%) | 48 (28%) | 30 (17%) |
| 4 | 11 | 44 | 2 (5%) | 34 (77%) | 5 (11%) | 3 (7%) |
| 5 | 1 | 5 | 0 (0%) | 1 (20%) | 1 (20%) | 3 (60%) |
| Total | 2792 | 3274 | 1574 (48%) | 832 (25%) | 580 (18%) | 288 (9%) |

We reported in Table 4 the quality results, for the different alternative splicing events, obtained by `ASGAL` and `SplAdder`— for the latter we used `Hisat2` and `STAR` as spliced aligner. More precisely, we computed the precision, recall, and F-measure values for the cases where we have only one event — distinguishing between exon skippings (ES), alternative acceptor (A3) and donor (A5) sites, and intron retentions (IR) — as well as complex scenarios composed of 2 to 5 events. Since the number of cases with more than one event is quite uncommon, we did not further refined these kind of events.

**Table 4.** Q uality measures in detecting alternative splicing events on the simulated datasets with 5M and 10M reads. Precision (Prec), recall (Rec), and F-Measure (FM) achieved on the simulated datasets in detecting alternative splicing events: exon skipping (ES), alternative acceptor site (A3), alternative donor site (A5), intron retention (IR), and genes in which more (from 2 up to 5) events were combined. Results obtained by `ASGAL` and `SplAdder`, using both `Hisat2` and `STAR` as spliced aligner, are reported.

| Sample | Event | ASGAL |  |  | SplAdder + Hisat2 |  |  | SplAdder + STAR |  |  |
| --- | --- | --- | --- | --- | --- | --- | --- | --- | --- | --- |
|  |  | Prec | Rec | FM | Prec | Rec | FM | Prec | Rec | FM |
| 5M | ES | 0.818 | 0.919 | 0.864 | 0.795 | 0.868 | 0.829 | 0.761 | 0.852 | 0.803 |
|  | A3 | 0.761 | 0.743 | 0.751 | 0.677 | 0.780 | 0.724 | 0.657 | 0.774 | 0.709 |
|  | A5 | 0.733 | 0.718 | 0.725 | 0.679 | 0.752 | 0.712 | 0.656 | 0.734 | 0.692 |
|  | IR | 0.645 | 0.666 | 0.654 | 0.370 | 0.483 | 0.417 | 0.395 | 0.483 | 0.432 |
|  | 2 | 0.848 | 0.721 | 0.778 | 0.800 | 0.670 | 0.729 | 0.737 | 0.656 | 0.693 |
|  | 3 | 0.895 | 0.540 | 0.673 | 0.761 | 0.459 | 0.572 | 0.824 | 0.459 | 0.589 |
|  | 4 | 0.760 | 0.431 | 0.549 | 0.720 | 0.409 | 0.520 | 0.680 | 0.386 | 0.491 |
|  | 5 | 1.000 | 0.800 | 0.888 | 1.000 | 0.800 | 0.888 | 1.000 | 0.800 | 0.888 |
| 10M | ES | 0.810 | 0.961 | 0.878 | 0.735 | 0.923 | 0.817 | 0.723 | 0.920 | 0.809 |
|  | A3 | 0.739 | 0.791 | 0.763 | 0.593 | 0.816 | 0.685 | 0.623 | 0.815 | 0.705 |
|  | A5 | 0.720 | 0.768 | 0.741 | 0.639 | 0.821 | 0.717 | 0.655 | 0.807 | 0.722 |
|  | IR | 0.664 | 0.673 | 0.667 | 0.357 | 0.542 | 0.429 | 0.390 | 0.549 | 0.455 |
|  | 2 | 0.840 | 0.762 | 0.799 | 0.693 | 0.662 | 0.676 | 0.725 | 0.661 | 0.691 |
|  | 3 | 0.838 | 0.568 | 0.675 | 0.745 | 0.505 | 0.601 | 0.738 | 0.534 | 0.619 |
|  | 4 | 0.730 | 0.431 | 0.540 | 0.692 | 0.409 | 0.514 | 0.750 | 0.409 | 0.528 |
|  | 5 | 1.000 | 0.800 | 0.888 | 1.000 | 0.600 | 0.750 | 1.000 | 0.600 | 0.750 |

The results show that `ASGAL` achieved the best values of precision, recall and F-measure in almost all the alternative splicing events with the only exception of the recall of the alternative splicing sites (A5 and A3). We investigated those cases and we found out that our method shows some limitations in detecting the alternative splicing site events that extend an annotated events: as previously described, to detect this kind of event, our method requires that the reads align leaving a gap on them, and requires the presence of sufficiently long anchors on two different exons. These requirements proved to be a bit too restrictive, and limited the capacity of our method to detect alternative splice site extending an exon.

However, our method achieved the best values of F-Measure in all the alternative splicing events, highlighting the ability of `ASGAL` in detecting the novel alternative splicing events. We note here that the row corresponding to cases with 4 AS events reports the worst values for `ASGAL`, while the same is not true for `SplAdder`. Therefore, we have investigated those cases and we observed that (and as shown in Table 3) the great majority (around 90%) of these events are alternative splicing sites (A5 and A3), which are the type of event achieving the worst results, as explained before. Moreover, as expected, by increasing the number of reads in the input set, both the tested methods achieve almost always better recall and worst precision.

Finally, for what concerns the efficiency of the two tested methods, the step of event identifications of `ASGAL` required ~ 2.7 seconds per annotation and 443MB of memory, while `SplAdder` required ~ 3.25 seconds and 47MB. Our approach used more memory since one of the steps consists in aligning the gaps left on the read against an intron, in order to detect the possible presence of alternative splice site events extending an annotated exon.

### Real Data

We also applied our method to a real dataset of RNA-Seq reads in order to assess its performance in detecting events from RNA-Seq data that are likely to be the result of the expression of a single transcript in each gene: this situation is the most common in real samples [29].

To do this, we considered the study proposed in [30], in which the role of the BRAF oncogene was investigated in melanoma cell migration. They considered the BRAFV600E mutation in melanoma skin cancer and on melanocytes overexpressing oncogenic BRAF to assess the effect on transcript expressions. The RNA-Seq experiments, conducted using the Illumina HiSeq 2000 sequencer, consisted of 2 BRAFv600e melanomas (SRR354042 and SRR354043), melanocytes+RFP control (SRR354040) and melanocytes + BRAFv600e (SRR354041) datasets of paired-end reads (GEO accession GSE33092). Since the goal of the presented analysis was to perform an in-depth analysis of the novel alternative splicing events induced by expressed transcripts, we decided to restrict ourselves only to genes with differentially expressed transcripts in the aforementioned datasets. In this way, we can ensure that all the transcripts we considered have a good support in terms of reads.

This was done by starting from the analysis done in [30], in which a list of differentially expressed transcripts of the BRAFv600e and the two melanoma datasets with respect to the control dataset. The list was obtained by running the pipeline presented in [31], in which Cufflinks [3] was applied to the spliced alignments computed by `TopHat2` [21]. Table 5 summarizes the datasets.

**Table 5.** Description of the real RNA-Seq datasets.

| SRA Accession | Condition | Num. Reads |
| --- | --- | --- |
| SRR354041 | Melanocytes + BRAFv600e | 100048222 |
| SRR354042 | Primary Melanoma 1 | 62884955 |
| SRR354043 | Primary Melanoma 2 | 71910612 |

As for the simulated dataset, we aligned the RNA-Seq reads of the three datasets to the reference genome with `STAR` [22] and we split each dataset into subsets corresponding to single genes based on read alignments, according to the human GENCODE annotation. Then, we selected only the subset of RNA-Seq reads corresponding to genes for which (at least) one transcript was found as differentially expressed in that dataset in [30]. This resulted in 630 genes for each dataset, which were used as input for `ASGAL`. As done in the simulated scenario, we built our ground truth using `AStalavista` extracting from its output all the alternative splicing events involving the transcripts of interests. This results in a total of 903 alternative splicing events: 366 exon skippings, 245 alternative acceptor sites, 156 alternative donor sites, and 136 intron retentions. Finally, for each gene, we generated a reduced annotation by removing the transcript found as differentially expressed. Using such reduced annotation, we run `ASGAL` and `SplAdder` comparing their results. Since we could not obtain any useful information from `SplAdder` when run with default parameters, we ran it providing different inputs to improve its results. First, we ran it setting the confidence level parameter to the smallest possible value to allow `SplAdder` to keep events with low support. Then, we used as input the alignments of the three samples simultaneously to provide a global view of the three samples and, finally, we merged the three alignments files into a single file and used it as input to check whether a single view of all the transcripts expressed could improve `SplAdder` results.

Table 6 shows the total number of alternative splicing events found by `ASGAL` and `SplAdder` in each of the considered RNA-Seq sample with respect to the reduced annotation while Table 7 shows the results of `SplAdder` obtained by considering the alignments of each sample simultaneously and by merging the three alignments files into a single file. The results obtained by our tool are similar to the results obtained with simulated data. Indeed, `ASGAL` is more effective in detecting exon skipping events than other splicing events, a common behaviour among tools for the detection of alternative splicing events [9], since exon skipping events are the easiest to detect. `SplAdder` achieved poor results compared to `ASGAL` in all the settings we tested, although the different settings improved its results. `SplAdder` obtains the best results when run providing a single input file containing all the alignments of all the sample; the main reason of this fact is that `SplAdder` is able to identify an event only if all the isoforms involved in it are supported by reads in the sample. This proves that `SplAdder`, at least when used without modifying the hard-coded parameters, is not suited to manage cases in which only one transcript is potentially expressed in the input RNA-Seq sample.

**Table 6.**
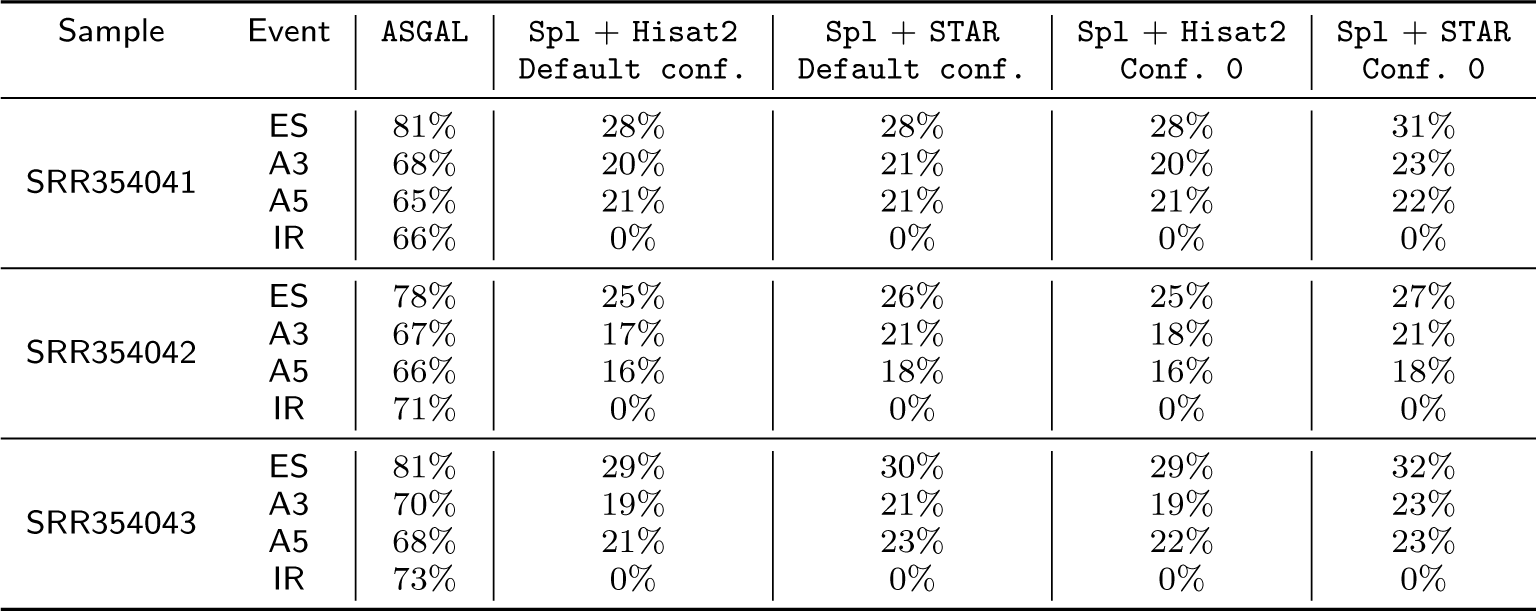
Number of AS events identified by `ASGAL` and `SplAdder` in the real data scenario with respect to the number of AS events identified by `AStalavista`. For each considered RNA-Seq sample and for each AS event type (ES: Exon Skipping, A3: Alternative Acceptor Site, A5: Alternative Donor Site, IR: Intron Retention), the ratio between these two numbers is shown. Results obtained by `ASGAL` and `SplAdder`, using both `Hisat2` and `STAR` as spliced aligner and setting the confidence level to the default value (3) and to the minimum one (0), are reported.

**Table 7.** Number of AS events identified by `SplAdder` in the real data scenario with respect to the number of AS events identified by `AStalavista`. For each AS event type (ES: Exon Skipping, A3: Alternative Acceptor Site, A5: Alternative Donor Site, IR: Intron Retention), the ratio between these two numbers is shown. Results obtained by `SplAdder`, using both `Hisat2` and `STAR` as spliced aligner, are reported. *Multiple sample* columns refer to the results obtained by `SplAdder` considering the alignments of each sample simultaneously while *Single sample* refer to the results obtained by merging the three alignments files into a single file.

| Event | Spl + Hisat2<br>Multiple sample | Spl + STAR<br>Multiple sample | Spl + Hisat2<br>Single sample | Spl + STAR<br>Single sample |
| --- | --- | --- | --- | --- |
| ES | 40% | 39% | 42% | 41% |
| A3 | 26% | 30% | 29% | 34% |
| A5 | 29% | 29% | 30% | 33% |
| IR | 0% | 0% | 0% | 0% |

Notice that `ASGAL` detected twice as many AS events as `AStalavista`. Since we could not measure the precision of `ASGAL` on real data, we verified that all the splicing events detected on real data were potentially expressed in the samples as follows. First, we extracted from the alignments computed with `STAR` all the identified introns, and then we compared them with the introns found by `ASGAL` used to detect the alternative splicing events. The results of this investigation show that 98% of the events identified by `ASGAL` but not by `AStalavista` are induced by an intron effectively expressed in the considered sample, confirming the novel alternative splicing events found by `ASGAL`.

## Conclusions

In this paper we propose `ASGAL`, a tool for predicting alternative splicing (AS) events from an RNA-Seq sample and a gene annotation given by a collection of annotated transcripts.

`ASGAL` differs from similar tools since it implements a splice-aware algorithm for mapping RNA-Seq data to a splicing graph. The alignments of the read to the splicing graph are then analyzed to detect differences, at the intron level, between the known annotation and the one obtained by the alignments, in order to reconstruct AS events.

Indeed, tools for AS prediction rely on a previously computed spliced-alignment of reads to a linear reference genome. While the spliced-alignment to a reference is a well understood notion, in this paper we investigate the problem of optimally mapping reads to a splicing graph by formalizing the notion of *spliced graph-alignment* and then propose an algorithmic approach to compute optimal spliced graph-alignments. Indeed, the graph aligner module of `ASGAL` can be used independently to produce spliced graph-alignments of RNA-seq reads to a general splicing graph.

Notice that our notion of spliced graph-alignment is tailored for detecting AS events that are either simple, or are a combination of two different simple events (see for example Figure 2, where the combination of an exon-skipping event with a competing event is represented). Such a notion deserves to be further investigated to detect more complex combinations of AS events. This will be the goal of a future development of the tool.

The experimental analysis discussed in this paper shows advantages in using a splice-aware aligner of RNA-seq data, and how it is useful in mitigating some recurrent problems that affect tools that detect AS events from differential expression of transcripts in multiple samples of RNA-seq data.

A problem which is related to mapping RNA-Seq reads to a splicing graph is mapping genomic reads directly to a graph representation of multiple genomes (pangenome): this problem is tackled by `vg` [32]. Despite this, how to apply a read mapper to a pan-genome graph for transcriptome analysis remains an interesting open problem.

## Ethics approval and consent to participate

Not applicable

## Consent for publication

Not applicable

## Availability of data and material

The datasets we used in the current study are publicly available at https://drive.google.com/open?id=1jnh5MS-B1eFqvu_ipvuL73NABv-hC1s9

## Author’s contributions

LD and RR developed the method, assisted by GDV and PB. PB, RR, and GDV designed the experimental setting. LD, SB, and MP implemented the method and performed the experimental analysis. PB supervised and coordinated the work. All the authors contributed to the manuscript writing.

## Acknowledgements

We acknowledge the support of the Cariplo Foundation grant 2013-0955 (Modulation of anti cancer immune response by regulatory non-coding RNAs).

## Footnotes

1 http://public.bmi.inf.ethz.ch/projects/2015/spladder/

2 Given the genomic region g of a gene, we compute the correct value as 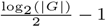 as described in `STAR` manual.

